# Intrinsic coupling modes reveal the functional architecture of cortico-tectal networks

**DOI:** 10.1101/014373

**Authors:** Iain Stitt, Edgar Galindo-Leon, Florian Pieper, Gerhard Engler, Eva Fiedler, Thomas Stieglitz, Andreas K Engel

## Abstract

In the absence of sensory stimulation or motor output, the brain exhibits complex spatiotemporal patterns of intrinsically generated neural activity. However, little is known about how such patterns of activity are correlated between cortical and subcortical brain areas. Here, we investigate the large-scale correlation structure of ongoing cortical and superior colliculus (SC) activity across multiple spatial and temporal scales. Cortico-tectal interaction was characterized by correlated fluctuations in the amplitude of delta, spindle, low gamma and high frequency oscillations (> 100 Hz). Of these identified coupling modes, topographical patterns of high frequency coupling were the most consistent with anatomical connectivity, and reflected synchronized spiking in cortico-tectal networks. Ongoing high frequency cortico-tectal coupling was temporally governed by the phase of slow cortical oscillations. Collectively, our findings show that cortico-tectal networks can be resolved through the correlation structure of ongoing neural activity, and demonstrate the rich information conveyed by high frequency electrocorticographic signals.

## Introduction

Rather than remaining entirely inactive in the absence of sensory stimuli or motor output, the brain displays complex spatiotemporal activation patterns that are more generally described as ‘ongoing activity’ (Arieli et al., 1996; Deco and Corbetta, 2010; Engel et al., 2013). For a long time ongoing activity was thought of as a form of neural noise, that was the net product of random fluctuations in neural networks. However, more recent studies have shown that the dynamics of spontaneously generated neural activity can be informative about the functional organization of large-scale brain networks (Fox et al., 2005; He et al., 2008; Hipp et al., 2012), revealing intrinsically generated coupling modes at multiple spatial and temporal scales (Deco and Corbetta, 2010; Engel et al., 2013). Converging evidence from both noninvasive and invasive approaches has led to the identification of the most prominent spectral features of large-scale ongoing functional interaction (Betti et al., 2013; Engel et al., 2013; Hipp et al., 2012; Phillips et al., 2014; Watrous et al., 2013). However for the most part, the study of ongoing brain dynamics has been heavily limited to measures of cortico-cortical functional coupling (He et al., 2008; Nir et al., 2008). Whether the result of an overly corticocentric focus of the field (Parvizi, 2009), or due to technical limitations of commonly employed experimental procedures, there have been relatively few studies on the role that subcortical structures play in the generation of intrinsically organized large-scale neural dynamics. It therefore remains unclear if the prominent coupling modes identified in cortex represent exclusively cortical phenomena, or if such coupling modes also extend to subcortical structures.

The superior colliculus (SC) presents itself an interesting model to study the ongoing dynamics of such cortical-subcortical functional interaction because it receives dense inputs from a wide range of sensory and motor cortical areas (Manger et al., 2010). In addition, the SC is highly connected with other subcortical structures that display reciprocal connectivity with cortex, such as the pulvinar and lateral geniculate nucleus (Abramson and Chalupa, 1988; Baldwin et al., 2011; Berman and Wurtz, 2011; Nakamura and Itoh, 2004). Therefore in the absence of sensory stimulation or motor output, the neural dynamics of the SC are presumably defined by the interaction of both cortical and sub-cortical inputs, and intrinsic SC network properties.

In this study, our aim was to use cortico-tectal networks in the anesthetized ferret as a model to study ongoing neural dynamics between cortical and subcortical structures. To obtain data spanning multiple spatial scales, we recorded spiking activity and local field potentials from all layers of the SC and visual cortex, while simultaneously sampling local field potentials (LFPs) from the entire posterior cortex using a custom-designed microelectrode array (μECoG, Figure 1). Using power envelope correlations (Hipp et al., 2012), we identified cortico-tectal functional coupling modes spanning several carrier frequencies. High frequency power envelope correlation was strongly related to the correlated spiking activity of SC and cortex neurons, and mirrored patterns of anatomical connectivity, while lower frequency coupling appeared less related to structural connectivity. Cortico-tectal functional interaction in the high frequency band was parcellated by the phase of slow cortical oscillations, reflecting the subcortical entrainment of ongoing activity to cortical ‘up’ and ‘down’ states. As the first demonstration of the large-scale correlation structure of ongoing cortico-tectal neural activity, these findings highlight that the functional architecture of large-scale networks in the brain can be resolved through the correlation analysis of ongoing neural dynamics.

**Figure 1.**
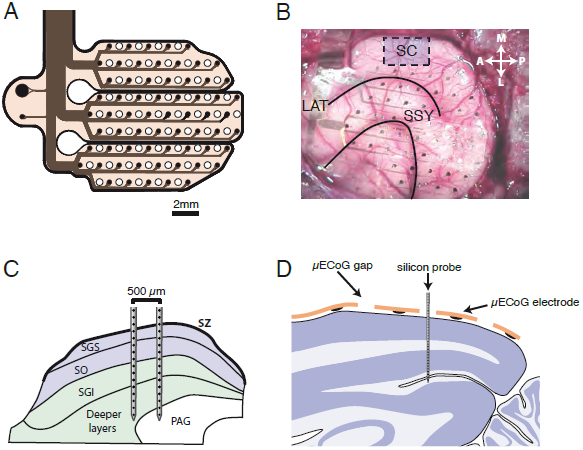
Experimental setup. (A) Schematic diagram of the custom designed μECoG array. 64 electrodes of 250μm diameter were distributed across 3 separate polyimide fingers and arranged in a hexagonal grid. Holes were cut into the polyimide foils in the space between electrodes to allow for the placement of linear silicon probes. (B) A photo from the surgical implantation of the μECoG array. The general area for SC penetrations is shown by a blue box. Black lines indicate the lateral sulcus (LAT) and the suprasylvian sulcus (SSY). (C) A schematic diagram illustrating the placement of dual-shank 32 channel silicon probes in the SC. Probes were placed such that neural data could be acquired from both superficial (blue) and deep (green) layers of the SC simultaneously. (D) A schematic illustration of the placement of linear silicon probes in the visual cortex. Single-shank 32 channel probes (100μm inter-electrode spacing) were advanced into the cortex through small holes in the liECoG array. Probes were advanced until the most superficial contacts were just above the pial surface, such that we recorded, in a single penetration, from superficial and deep visual cortex simultaneously. Abbreviations: SZ, *stratum zonale;* SGS, *stratum griseum superficiale;* SO, *stratum opticum*; SGI, *stratum griseum intermediale*; PAG, periaqueductal gray; ICP, pericentral nucleus of the inferior colliculus; ICX, external nucleus of the inferior colliculus; ICC, central nucleus of the inferior colliculus.

## Results

### Large-scale correlation structure of cortico-cortico and cortico-tectal interactions

Before assessing the dynamics of simultaneously recorded SC and cortical activity, we first wanted to identify the spectral signatures that define cortico-cortico functional connectivity under isoflurane anesthesia (for average ongoing power spectra of SC, intracortical and μECoG recording sites, see Supplementary Figure 1). To this end, we computed the correlation of band-limited signal power envelopes between all possible combinations of μECoG recording contacts. Figure 2A illustrates the strength of power correlation for each carrier frequency as a function of μECoG inter-electrode distance. Power envelopes of oscillations in the slow (∼0.7 Hz), delta (∼3 Hz) and spindle (∼11 Hz) frequencies were correlated over large distances in the cortex. For frequencies above 30 Hz, power envelope correlation gradually decreased with increasing frequency such that μECoG signal frequencies above 120 Hz displayed minimal inter-electrode power envelope correlation, suggesting that such high frequency μECoG signal components reflect neural activity at a more local scale. To validate cortico-cortico functional connectivity findings from μECoG LFPs, we repeated the same power envelope correlation analysis between intracortical recording sites and μECoG contacts (Figure 2A). Indeed, intracortical-μECoG power envelope correlation displayed the same spectral characteristics, with spontaneous power fluctuations strongly correlated for slow (∼0.7 Hz), delta (∼3 Hz) and spindle (∼11 Hz) frequencies over larger distances.

**Figure 2.**
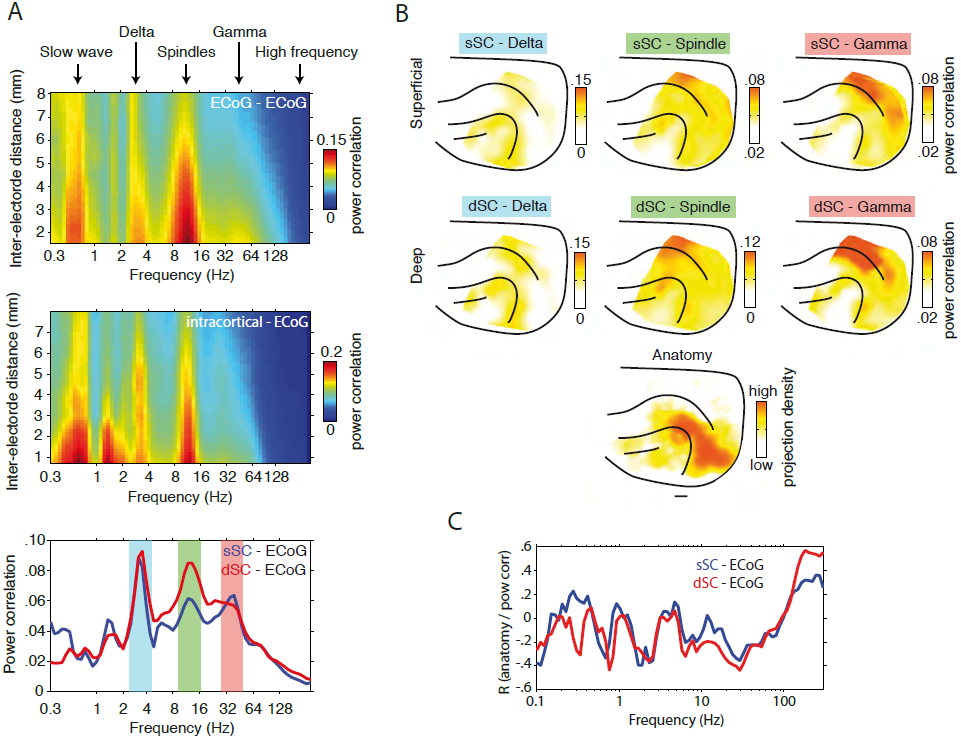
Dynamics and large-scale topography of cortico-cortical and cortico-tectal power envelope correlation. (A) Top: Population-averaged power envelope correlation computed between μECoG contact pairs separated by varying distances. Note that correlations are widely distributed in slow (∼0.7 Hz), delta (∼3 Hz) and spindle (8-15 Hz) frequency ranges and that power envelopes are minimally correlated for frequencies above 120 Hz. Middle: population averaged power envelope correlation computed between intracortical and μECoG recording sites. Bottom: population averaged cortico-tectal power envelope correlation computed for SC recording contacts located in superficial (blue) and deep (red) SC layers. Note the peaks in cortico-tectal power correlation for delta, spindle and gamma frequencies. (B) The average cortical topography of cortico-tectal power correlation for delta, spindle and gamma carrier frequencies. Maps are plotted for both superficial and deep SC layers. To compare functional coupling to anatomy, we plotted the density of tectally projecting neurons across the cortical surface. Anatomical data were adapted with permission from Fig. 1 in Manger et al (2010). (C) Spatial correlation of anatomical connectivity (B, bottom) and functional coupling topographies across all frequencies. Analyses for superficial and deep layers are plotted in blue and red respectively.

Since the origin of cortical afferent inputs varies between superficial and deep SC layers (Harting et al., 1992), we used current source density analysis to separate SC recording contacts into superficial and deep zones (Supplementary Figure 2) for cortico-tectal functional connectivity analysis (Stitt et al., 2013). Reflecting cortico-cortico power correlation spectra, global cortico-tectal power envelope correlation was characterized by peaks in the delta, spindle, and low gamma frequencies (Figure 2A, bottom). Cortico-tectal coupling in the delta frequency displayed a patchy cortical topography, while spindle coupling was widespread with a peak in parietal cortical areas (Figure 2B). Coupling in the low gamma frequency displayed a more spatially specific cortical topography, with a clear peak in parietal areas (Figure 2B). Given that anatomical connectivity defines the structural framework upon which neural dynamics is superimposed, functional connectivity measures reflecting direct corticotectal interaction should overlap with patterns of anatomical connectivity. To test this, we computed the spatial correlation of cortico-tectal coupling topographies with the distribution of SC-projecting neurons across the cortex (Figure 2C). Cortico-tectal anatomical connectivity data were adapted with permission from Manger et al (2010). None of the identified delta (sSC r = -0.11; sDC r = -0.12), spindle (sSC r = -0.09; dSC r = -0.22) or gamma (sSC r = -0.27; dSC r = -0.33) frequency bands displayed coupling profiles that matched anatomical connectivity patterns. However, despite displaying quantitatively weaker power envelope correlation, coupling in frequencies above 100 Hz mirrored anatomical connectivity for both superficial (r = 0.29) and deep (r = 0.47) SC layers (Figure 2C). These results suggest that coupling in the identified delta, spindle, and gamma frequency bands reflect indirect modes of cortico-tectal functional interaction, whereas coupling in high frequencies likely reveals direct cortico-tectal communication.

### Correlated fluctuations in high frequency LFPs reflect cortico-tectal structural connectivity

We next investigated the dynamic relationship between simultaneously recorded high frequency (> 120 Hz) LFP components in the SC and cortex. Figure 3A displays a short time-series of co-recorded SC, intracortical, and μECoG data from an example experiment. Power envelopes of high frequency signals displayed spontaneous bursting-like activity patterns that appeared to occur synchronously between the SC and cortex. To visualize the cortical topography of SC-μECoG interactions, we selected a seed channel in the SC and plotted the strength of power correlation with all μECoG recording contacts as a heat map on a model ferret brain (Figure 3B). The example SC recording contact shown in Figure 3 displays spatially specific power correlation with two separate clusters of μECoG channels over lateral visual cortex, and the suprasylvian gyrus. Cortico-tectal power correlation topographies measured using high frequencies were consistent when computed across non-overlapping time periods (Supplementary Figure 4). In contrast to the SC, high frequency signals that are recorded intracortically only display power envelope correlation with immediately surrounding μECoG electrodes in the lateral visual cortex (Figure 3C). In this example recording, the cortical topography of SC-μECoG, and intracortical-μECoG power correlation display considerable overlap in the lateral visual cortex (lower blob in Figure 3B and 3C). Reflecting this overlap, we also observed power envelope correlation between a cluster of SC and intracortical channel pairs (Figure 3D).

**Figure 3.**
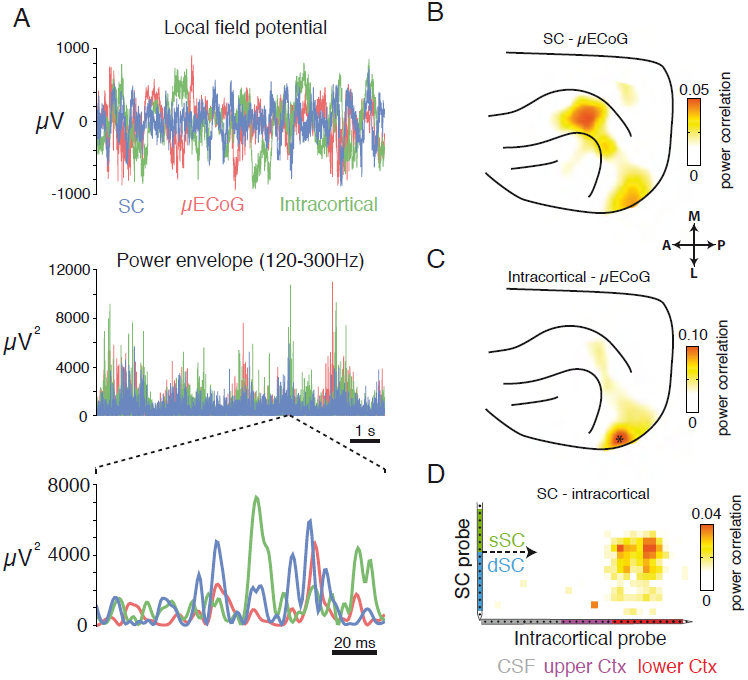
High frequency LFP power envelopes are correlated between cortex and SC. (A) Simultaneously recorded SC, intracortical, and μECoG signals from one example recording session. No relationship between signals is immediately visible upon inspection of LFPs (top). However, high frequency power envelopes display burst-like fluctuations that appear correlated at both slow (middle) and fast (bottom) timescales. (B) The cortical topography of high frequency power envelope correlation for the seed SC electrode displayed in A. Note the region-specific power correlation for lateral visual and suprasylvian cortical areas. (C) The cortical topography of high frequency power envelope correlation for the intracortical recording site displayed in A. Note the presence of correlated high frequency activity in cortical regions immediately surrounding the position of the intracortical recording site (marked as ^*^). (D) Displays the strength of high frequency power correlation between all SC and intracortical channel pairs. Note the presence of a cluster of correlated channels in the center of the SC probe, and the lower third of the intracortical probe.

We next computed the correlation of high frequency LFP power envelopes between all possible combinations of SC-μECoG and SC-intracortical recording sites. Figure 4 displays the average cortical topography of high frequency LFP power correlations from superficial SC to μECoG (Figure 4A, left), and deep SC to μECoG (Figure 4A, middle). Superficial SC recording sites were correlated with μECoG contacts distributed over the entire visual cortex, with strongest correlation in cortical area 18 (power correlation = 0.019 ± 0.005 SEM), slightly weaker correlation in higher visual (SSY: Suprasylvian area, 0.014 ± 0.004) and PPc (0.012 ± 0.003) areas, and almost no correlation in auditory cortical areas (0.005 ± 0.002) (Figure 4B, left). In contrast, deep SC layers displayed power correlation effects with a wider range of cortical areas, with strongest correlation to visual area 21 (0.021 ± 0.005 SEM) and posterior parietal areas (PPc = 0.021 ± 0.004, PPr = 0.020 ± 0.005) (Figure 4B, middle). The cortical topography of deep SC to μECoG high frequency correlation extended from early visual areas, through higher visual and multisensory areas along the suprasylvian gyrus towards somatosensory cortex, and reflecting the data presented in Figure 2C, displayed a striking similarity to the topography of cortico-tectal anatomical connectivity (Figure 4A, right).

**Figure 4.**
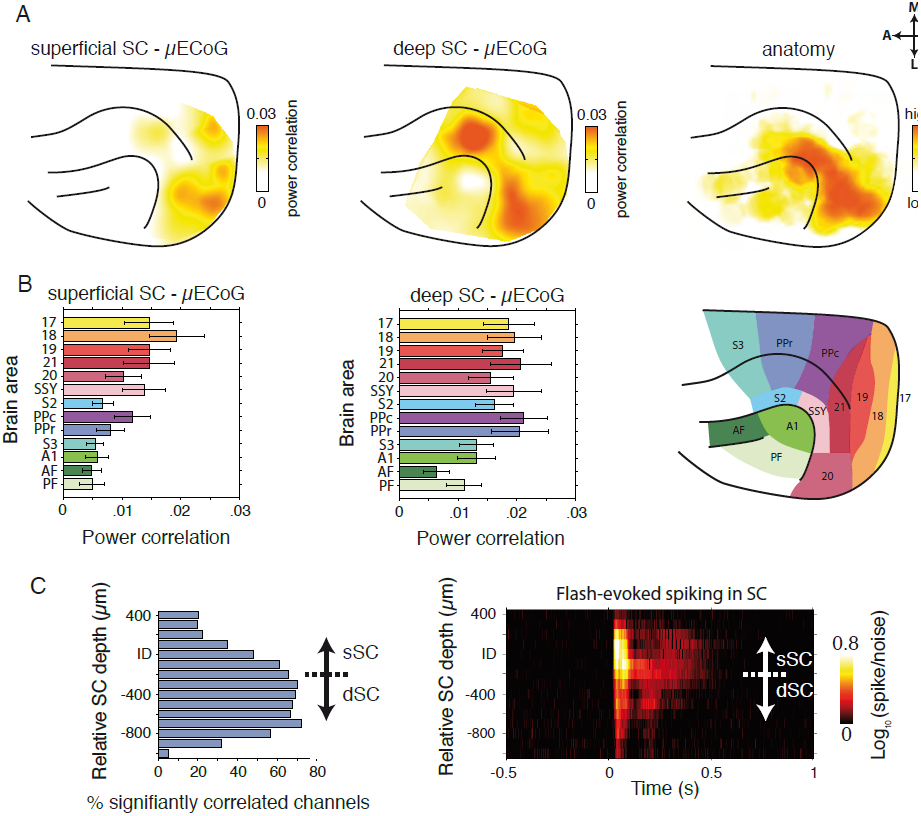
Large-scale topography of high frequency cortico-tectal power envelope correlation. (A) Left: The average cortical topography of LFP power correlation between superficial SC andarray. Note the strongest power correlation over posterior visual cortex. Middle: The average cortical topography of deep SC to μECoG LFP power correlation. Note the presence of strong power correlation in visual and suprasylvian cortical areas. Power correlation effects extend anterior-medially along the suprasylvian gyrus towards a separate cluster of strong correlation in suprasylvian cortex. Right: Plot of the density of tectally projecting neurons across the cortical surface. Data were adapted with permission from Manger et al (2010). (B) Population averaged (± SEM) strength of superficial SCμECoG (left) and deep SCμECoG (middle) high frequency power envelope correlation for different cortical areas. A map of the areal parcellation used in this analysis is shown on the right. Note the presence of strong correlation in early visual cortical areas for superficial SCμECoG channel pairs. In contrast, deep SCμECoG power correlation was more widespread, encompassing visual, suprasylvian and posterior parietal areas. (C) Left: Incidence of significant SCμECoG high frequency power correlation as a function of SC depth. Right: Population averaged spike/noise ratio in response to visual flash stimulation (flash onset at 0 s). Note that significant power correlation with cortex was most prominent in intermediate/deep SC layers.

To assess cortico-tectal power correlation effects systematically across SC layers, we aligned all penetrations to the current source density inflection depth (see Supplementary Figure 2) and then computed the percentage of recording sites that were significantly correlated with cortex at each depth (Figure 4C). Cortico-tectal power correlation effects displayed a clear depth dependency in the SC, with correlation being lower in upper superficial layers, gradually increasing with depth to peak in intermediate layers, before fading again in the deepest layers (for cortical depth profile, see Supplementary Figure 5).

### High frequency LFP power correlation reflects correlated cortico-tectal spiking activity

We reasoned that correlated fluctuations of high frequency extracellular fields between SC and cortex might to a large extent reflect the synchronous spiking activity of neurons across cortico-tectal networks. To test if SC spiking occurred synchronously with fluctuations in high frequency μECoG power, we computed SC spike triggered average (STA) spectrograms based on μECoG signals. Figure 5A displays an example recording, with SC STA spectra shown for μECoG contacts that displayed both weak and strong high frequency power envelope correlation respectively. There was little observable change in μECoG signal power locked to the timing of SC spikes for the channel pair that displayed weak power correlation. In contrast, the SC-μECoG channel pair that displayed strong power correlation showed a broadband increase in the power of μECoG frequencies above 10 Hz locked to the timing of SC spiking activity (Figure 5A). Generalizing from the example shown in Figure 5A, SC-μECoG channel pairs displaying significant high frequency coupling showed increased STA μECoG power in frequencies above 10 Hz locked to the timing of SC spikes (Figure 5B). In addition, the magnitude of cortico-tectal power correlation was tightly coupled to the STA power of high frequency cortical signals across all SC-μECoG channel pairs (r = 0.68, p < 0.0001), indicating that spontaneous fluctuations of high frequency μECoG signal power occur synchronously with spiking activity in the SC (Figure 5C).

**Figure 5.**
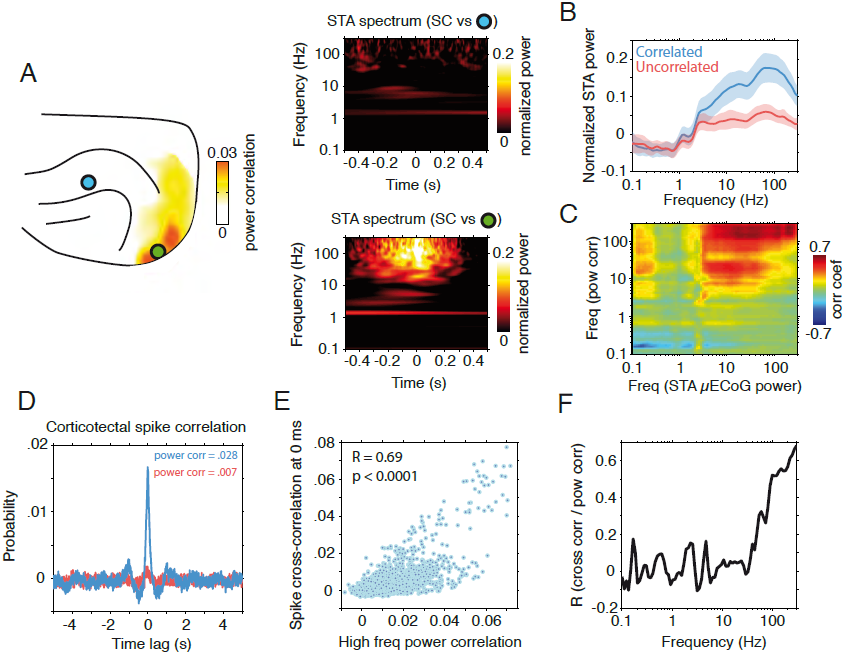
High frequency power envelope correlation reflects synchronized cortico-tectal spiking activity. (A) SC spike triggered average spectrograms for two different μECoG contacts from an example recording session. The location of the contacts is shown in the panel on the left. Both spectrograms were computed using spiking data from the same SC seed contact. The strength of cortico-tectal power correlation for the SC seed electrode is displayed as a heat map. Note the presence of an increase of μECoG signal power for frequencies above 20 Hz timelocked to SC spiking activity for the channel pair that showed strong power correlation (right, bottom panel). In contrast, the weakly correlated channel pair displayed no spike-triggered change in signal power (right, top panel). (B) The population mean (± standard deviation) for the spike triggered average spectrograms. SC-μECoG channel pairs displaying significant high frequency power correlation are plotted in blue, while all others are plotted in red. (C) The correlation of power envelope coupling and spike triggered average power for all frequency-frequency combinations. Note that high frequency coupling is correlated to the strength of spike triggered average signals in frequencies above ∼3 Hz. (D) Example spike cross-correlation histograms for SC-intracortical channel pairs that displayed strong (blue) and weak (red) high frequency power envelope correlation respectively. Note the large peak around ∼0 ms for the correlated channel pair, and the lack of structure for the uncorrelated channel pair. (E) Scatter plot of the strength of power correlation plotted against the probability of synchronous spiking for all SC-intracortical channel pairs. The correlation coefficient and related p-value are shown as an inset. (F) Correlation of power envelope coupling and synchronized spiking activity as a function of frequency. Note that coupling in high LFP frequencies specifically correlates with synchronized cortico-tectal spiking activity.

To confirm the neurophysiological origin of high frequency cortico-tectal power correlation, we next investigated the relationship between LFP power envelope correlation and cortico-tectal spike-spike correlations. Figure 5D displays spike cross-correlation histograms for two SC-intracortical channel pairs that display strong and weak power correlation respectively. The correlated channel pair shows a large peak centered at 0 ms in the cross-correlation histogram, indicating that these neurons fire synchronously. In contrast, the uncorrelated channel pair showed little or no structure in the spike cross-correlation histogram, indicating these co-recorded neurons fire independently. Across all SC-intracortical channel pairs, the strength of high frequency power correlation was significantly correlated with the probability of synchronized cortico-tectal spiking (r = 0.69, p < 0.0001, Figure 5E). To test if this effect was specific for high frequency coupling, we repeated the same analysis across all frequencies (Figure 5F). Indeed, this confirms that high frequency power envelope correlation specifically tracks the synchronized spiking activity of co-recorded neuron populations, with coupling at frequencies below approximately 60 Hz displaying weak correlation with coordinated spiking activity.

### High frequency LFP power correlation during sustained visual stimulation

To investigate how patterns of cortico-tectal LFP power correlation change following the application of an external driving force (i.e., sensory stimulation), we performed power correlation analysis on neural activity induced by drifting grating stimulation. To ensure that we analyzed only sustained activity throughout stimulation, high frequency power correlation was computed in the time window from 100 ms after grating onset, up until the offset of stimulation. In the SC, drifting gratings typically evoked large power increases in frequencies above 80Hz (Figure 6A), while μECoG responses displayed power increases in a broader range of frequencies above 40 Hz (Figure 6A). Across all SC-μECoG electrode pairs there was a strong correlation between the strength of spontaneous and stimulus driven high frequency (> 120Hz) power correlation (r = 0.45, p < 0.001), indicating that patterns of cortico-tectal functional connectivity were generally preserved across spontaneous and stimulus driven conditions. However, the strength of power correlation during visual stimulation was significantly weaker than spontaneous correlation (Figure 6A, number of data points below 1:1 dotted line, p < 0.001, sign test). This result suggests that dynamic fluctuations of SC neural activity are more dependent on cortical inputs during ongoing activity, while bottom up retinal inputs and intrinsic network properties may play a larger role in shaping the dynamics of SC activity during visual stimulation.

**Figure 6.**
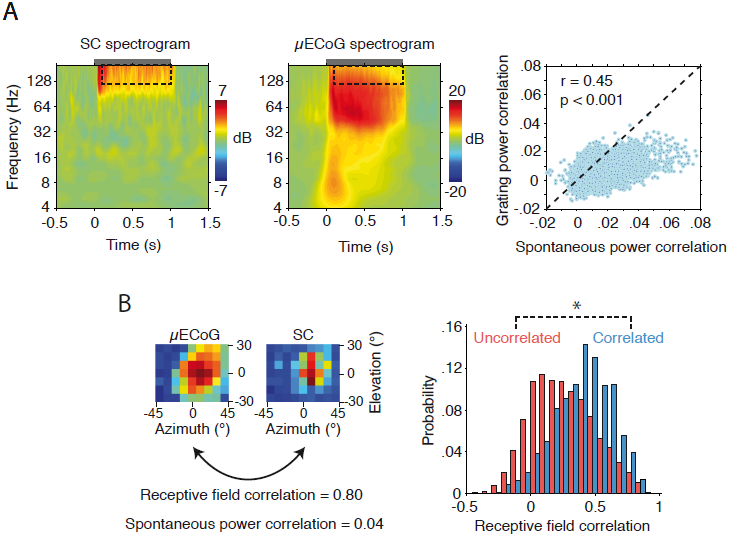
Comparison of spontaneous and stimulus induced high frequency neural dynamics. (A) Drifting grating induced power changes for representative SC (left) and μECoG (middle) recording sites. The gray bar above each spectrogram indicates the duration of drifting grating stimulation. Note the broadband increase in high frequency power for both SC and μECoG recording sites. Right: A scatter plot of the strength of spontaneous SC-μECoG verses drifting grating induced SC-μECoG high frequency power correlation. Note that most data points lie below the 1:1 line, indicating that SC-μECoG channel pairs display stronger power correlation during spontaneous activity. (B) Left: μECoG and SC visual spatial receptive fields from one example recording session. Note that this SC-μECoG channel pair displayed considerable visual spatial receptive field overlap as well as strong high frequency power envelope correlation during spontaneous activity. Right: Histogram displaying the distribution of visual receptive field correlation for all spontaneously power correlated and uncorrelated SC-μECoG channel pairs. ^*^ p < 0.01

### Correlated cortico-tectal recording sites display similar visual spatial receptive fields

We next tested if SC-μECoG channel pairs identified through high frequency LFP power correlation analysis displayed similar visual spatial receptive fields. Since previous anatomical studies have shown that visual cortical neurons project to areas of the SC with corresponding retinotopic representations (Berson, 1988), we reasoned that spontaneously correlated corticotectal recording sites should display similar spatial receptive fields. Figure 6B displays SC and μECoG visual spatial receptive fields from an example channel pair that exhibited significant high frequency power envelope correlation. Despite the SC receptive field being smaller than the μECoG receptive field, both recording sites responded to stimuli in similar locations in the visual field. To quantify the similarity of SC and μECoG spatial receptive fields, we computed the correlation coefficient between the two-dimensional visual spatial response matrices (Figure 6B). Spontaneously coupled SC-μECoG channel pairs showed significantly stronger similarity of visual spatial receptive fields (Power correlated: mean RF correlation = 0.46 ± 0.22 SD, power uncorrelated: mean RF correlation = 0.27 ± 0.25 SD, p < 0.0001, Figure 6B). These results confirm that functionally coupled cortico-tectal recording sites display similar visual receptive fields, as predicted by anatomical connectivity.

### Correlated cortico-tectal activity is temporally coordinated by slow oscillations

To further investigate the temporal dynamics of cortico-tectal functional connectivity, we computed the phase-locking value (PLV) of SC spiking activity to cortical oscillations. In general, SC spiking activity was strongly modulated by the phase of cortical oscillations at a frequency of approximately 0.8 Hz (Figure 7A, left), matching the reported frequency of the slow cortical oscillation (Crunelli and Hughes, 2010; Steriade et al., 1993a). However, functionally coupled SC-μECoG channel pairs displayed significantly stronger phase-locking to slow cortical oscillations than uncorrelated channel pairs (Correlated: PLV = 0.09 ± 0.002 SEM, Uncorrelated: PLV = 0.05 ± 0.001, p < 0.01). Cortico-tectal slow oscillatory spike-phase locking was specific for endogenous cortical slow oscillations, and displayed no temporal dependency to the frequency of artificial ventilation (Supplementary Figure 6). In contrast to corticotectal spike-phase locking, SC spiking activity displayed significant phase locking to local oscillations at a frequency of approximately 10Hz (PLV = 0.055 ± 0.002 SEM, p < 0.01) (Figure 7A, right).

**Figure 7.**
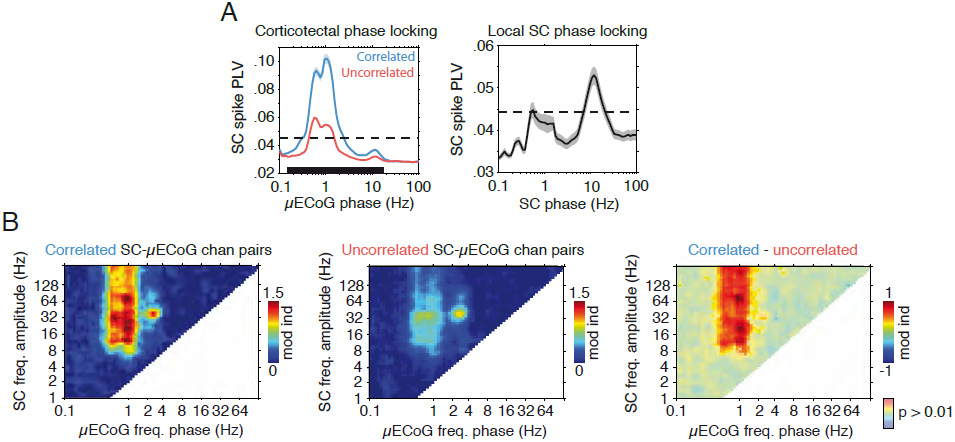
The dynamics of spontaneous SC neural activity are dominated by the phase of slow and spindle oscillations. (A) Population averaged (± SEM) SC spike phase locking values calculated using the phase of cortical (left) and local SC oscillations (right). The dashed line in each plot illustrates the level of significance (p < 0.01). For SC-μECoG channel pairs, significantly correlated and uncorrelated channel pairs are plotted separately. The black bar at the bottom of the SC-μECoG plot indicates the area in which correlated and uncorrelated curves are significantly different (p < 0.01). Note that spiking in power correlated channel pairs is more strongly locked to the phase of slow cortical oscillations. In addition, SC spiking activity is additionally locked to the phase of local oscillations at the spindle frequency (∼10Hz). (B) Population averaged cross-frequency phase-amplitude spectrograms calculated using the phase of μECoG oscillations and the amplitude of SC signals. Significantly correlated SC-μECoG channel pairs are shown on the left, uncorrelated channel pairs in the middle, and the difference between correlated and uncorrelated on the right. Note that SC-μECoG channel pairs that display strong high frequency power correlation also show significantly stronger coupling of SC oscillations above 8 Hz to the phase of slow cortical oscillations.

To quantify the relationship between SC and cortical oscillations by an additional coupling measure that takes phase into account, we computed the cross-frequency phaseamplitude coupling between SC and μECoG recording sites (see Supplementary Figure 7). This measure quantifies the strength with which the amplitude of SC LFP oscillations is modulated by the phase of μECoG signals across all possible frequency-frequency combinations. Figure 7B displays population averaged cross-frequency phase-amplitude spectrograms for SC-μECoG channel pairs that display significant (left) and insignificant (middle) high frequency power correlation respectively. Cross-frequency coupling for correlated SC-μECoG channel pairs was characterized by the strong modulation of SC activity above 8 Hz by the phase of slow cortical oscillations. In contrast, uncorrelated SC-μECoG channel pairs displayed comparatively weak cross-frequency coupling (Figure 7B, middle). The difference in cross-frequency spectra between significantly correlated and uncorrelated SC-μECoG channel pairs reveals that the phase of slow cortical oscillations specifically modulate the amplitude of SC oscillations for functionally coupled cortico-tectal recording sites (Figure 7B, right, p < 0.01, Bonferroni corrected).

To illustrate the dependency of SC activity on cortical slow oscillation phase in raw data, Figure 8A displays a raster plot of spontaneous SC spiking activity plotted with a simultaneously recorded μECoG signal bandpass filtered in the slow oscillatory frequency range (0.5 – 1.2 Hz). In this short epoch, SC spiking activity across several layers appears to group on the downward phase of the slow cortical oscillation. Despite most channels being locked to the phase of the slow oscillation, closer visual inspection of the raster plot reveals that recording sites at different depths within the SC are locked to slightly different phases of the cortical oscillation. Indeed, spike-phase histograms shown in Figure 8B exhibit slightly different distributions, with spiking activity from the more superficial recording contact centered on a slightly later μECoG slow oscillation phase than the deeper recording contact. We next quantified the preference of SC spiking activity to slow cortical oscillation phase by computing spike-phase histograms across all SC layers. Spiking activity across all SC layers showed a preference for the slow oscillatory phase corresponding to cortical ‘up’ states; with the strongest phase dependency detected in intermediate SC layers, 300μm below the presumed superficial/deep border (Figure 8C). In addition to displaying stronger modulation by slow cortical oscillations, spiking activity in intermediate layers also occurred at an earlier phase than recording sites located both dorsally in superficial layers, and ventrally in the deep SC (relative phase at distance to inflection depth; 16° at +200μm, -22° at -500μm and -11° at -1000μm). These findings demonstrate that while all SC layers are entrained during cortical ‘up’ states, intermediate SC layers are entrained first, before activity gradually propagates in both superficial and deep directions.

**Figure 8.**
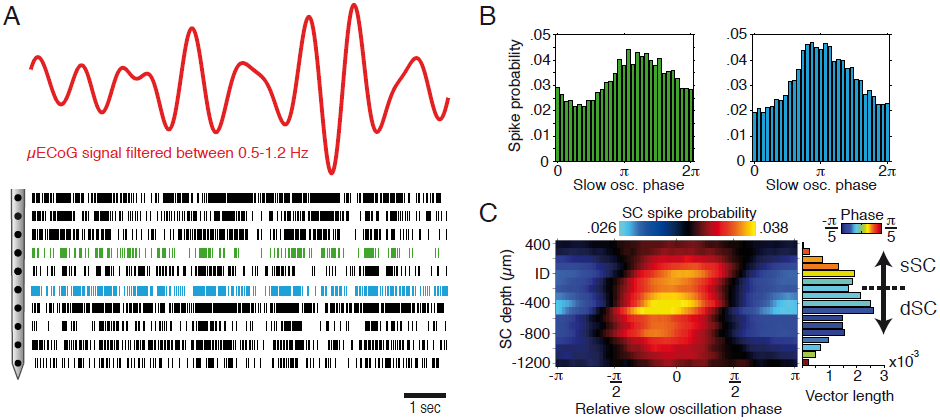
The dependency of SC spiking activity to the phase of cortical slow oscillations. (A) Example of μECoG signal filtered at the slow oscillation frequency with a raster plot containing spiking data from several SC channels. Note that the bursting behavior of SC neurons appears to be phase locked to the trough of the filtered slow wave. (B) Spike-phase histograms for the two channels marked (blue and green) in A. Note that neurons from the more superficial recording contact prefer a slightly earlier phase than the deeper recording contact. (C) Population averaged spike-phase histogram as a function of SC depth. Histograms were compiled using SC spiking activity and the phase of μECoG slow oscillations, and centered on the slow oscillatory phase corresponding to cortical ‘up’ states , where a relative phase of 0 indicates the center of the ‘up’ state. The strength with which average spike-phase histograms across each SC depth deviate from a uniform circular distribution is shown as a bar plot to the right, where the color of each bar indicates the preferred phase at each depth. Note that spike-phase locking is strongest, and occurs at the earliest phase in intermediate SC layers. From intermediate layers, both the strength of phase locking and phase lag decrease gradually with increasing distance in dorsal and ventral directions.

## Discussion

To summarize, this study reveals for the first time the complex spatiotemporal structure of correlated cortico-tectal neural activity. We identified several distinct cortico-tectal coupling modes spanning different carrier frequencies, with power envelope correlation in low gamma and high frequencies likely reflecting modes of indirect and direct communication respectively. Patterns of high frequency power correlation specifically matched the topography of anatomical connectivity. We demonstrated that such coincident fluctuations in high frequency LFPs tracked the synchronized spiking activity of SC and cortical neurons. Correlated cortico-tectal neural activity was temporally governed by the phase of slow cortical oscillations, reflecting the subcortical propagation of cortically generated ‘up’ states. Together, these findings reveal the multiple temporal and spatial scales of cortico-tectal functional interaction, and the highlight rich information content of high frequency LFPs measured by electrocorticogram.

### Cortico-tectal coupling modes

Reflecting the multiple temporal scales of cortico-cortico functional interaction, our results confirm the existence of several cortico-tectal coupling modes with distinct carrier frequencies. From these identified coupling modes, power envelope correlation in the low gamma and high frequencies displayed the most spatially specific cortico-tectal interaction, with peaks in parietal and visual/suprasylvian cortical areas respectively. Indeed, reflecting the inherent link between structural connectivity and functional interaction, we found that the spatial topography of cortico-tectal high frequency power correlation is in strong agreement with previously identified patterns of anatomical connectivity (Manger et al., 2010), with visual areas 18, 21, and medial suprasylvian areas providing the densest cortical input in the ferret (Harting et al., 1992). In contrast to high frequencies, functional interaction in delta, spindle, and low gamma frequencies displayed spatial topographies that were inconsistent with direct cortico-tectal anatomical connectivity. Since the SC is highly interconnected with other subcortical structures such as the substantia nigra (Harting et al., 1988; Kemel et al., 1988), pulvinar (Abramson and Chalupa, 1988; Baldwin et al., 2011; Berman and Wurtz, 2011), ventral lateral geniculate nucleus (Conley and Friedekich-ecsy, 1993; Nakamura and Itoh, 2004), and lateral posterior nucleus (Lugo-garcia and Kicliter, 1987), it is likely that coupling measured in these frequency bands reflects indirect cortico-tectal interaction modulated by other factors. Or alternatively, such coupling may reflect a more general, brain-wide form of coordinated activity, similar to resting-state networks identified in human studies (Hipp et al., 2012). Nevertheless, the disparity between low frequency and high frequency coupling modes reveals that independent and complementary information about functional interaction is conveyed through different physiological carrier frequencies. In general, our results are compatible with the hypothesis that cortico-cortico and cortico-tectal coupling modes operate under the same unifying neurophysiological principles. This finding is similar to the results of Saalmann et al (2012), who found that long-range synchronization between V4 and the tempero-occipital area (TEO) in macaque cortex was coordinated by the pulvinar. However unlike the pulvinar, which coordinates cortical activity, the SC – by virtue of its unidirectional input from the cortex – reads out cortical activity. Collectively, these findings are approaching a unified framework of interregional functional interactions in the brain, where subcortical structures act in unison with cortex to produce dynamic spatiotemporal patterns of activity. Although commonly identified coupling modes are present in the cortex, one cannot discount the role of subcortical structures, either as a driving force (i.e., pulvinar) or reading out activity (i.e., SC).

### Resolving networks by analysis of ongoing activity

To our knowledge, this study represents the first attempt at investigating the large-scale spatiotemporal structure of spontaneous cortico-tectal interaction, therefore to place our results into context we must look to previous studies investigating different brain areas. Our results are most similar to a study performed by Fukushima and colleagues (2012), which showed that high frequency μECoG signals in awake monkey auditory cortex display spatial covariations in a manner that is reflective of the underlying tonotopic map of the auditory cortex. Revealing a similar functional organization of the primary visual cortex of the cat, Kenet and coworkers (2003) showed using voltage imaging that the spatiotemporal structure of spontaneous neural activity was highly correlated to the underlying orientation map. Aside from reflecting the functional organization or brain regions, large-scale spatiotemporal fluctuations of spontaneous neural activity have also been shown to explain a large degree of trial-to-trial variability of sensory evoked responses in primary visual cortex (Arieli et al., 1996). However, unlike the abovementioned studies, which report the correlation structure of indirect measures of neural activity (i.e., surface potentials and voltage-sensitive dye), we report here a direct link between the spontaneous correlation structure of high frequency extracellular fields and the synchronized spiking activity of widely distributed neurons. Similar to our findings, Nir and coworkers (2008) showed that time-to-time variations in neuronal firing rate are strongly correlated to power modulations of high frequency extracellular fields (40-100Hz) between interhemispheric recording sites in the human auditory cortex. Other groups have investigated the correlation structure of intrinsically generated brain activity using functional magnetic resonance imaging (fMRI), which tracks the slow modulations of blood oxygen level dependent (BOLD) signals throughout the brain (Hutchison et al., 2013). Similar to our findings, Vincent and coworkers (2007) showed that spontaneous BOLD fluctuations in the anesthetized monkey brain are correlated between anatomically connected cortical regions. Since BOLD signals track slower hemodynamic responses of brain regions (an indirect measure of neural activity), taken together, these studies reveal that functional brain organization can be delineated at both very slow and very fast temporal scales. Collectively, power correlations in neural activity measured electrophysiologically or through fMRI can be more generally described as envelope intrinsic coupling modes (Engel et al., 2013). Although only demonstrated in cortico-tectal networks here, we propose the correlation analysis of spontaneous high frequency power envelopes as a useful tool for elucidating direct functional connectivity in high throughput invasive recording paradigms.

### SC activity couples to cortical slow and thalamic spindle oscillations

As shown here, the cortical slow oscillation temporally regulates correlated corticotectal activity under anesthesia. Slow cortical oscillations are generated locally within cortical layer 5, and subsequently propagate to other granular and supragranular cortical layers (Sanchez-Vives and McCormick, 2000; Steriade et al., 1993a, 1993b). Under normal physiological circumstances, slow cortical oscillations are the defining feature of slow wave sleep (Mölle and Born, 2011), however, they are also present under anesthesia, with the emergence of slow cortical oscillations representing the strongest physiological correlate of the loss of consciousness following the induction of propofol anesthesia (Lewis et al., 2012). Since projection neurons in cortical layer 5 are strongly activated during up states of the slow oscillation (Contreras and Steriade, 1995; Cowan and Wilson, 1994; Sanchez-Vives and McCormick, 2000), it is perhaps unsurprising that the downstream target neurons of these cells fire preferentially during cortical up states. Previous studies have also shown that neural activity in the basal ganglia (Magill et al., 2000), thalamus (Timofeev and Steriade, 1996), cerebellum (Ros et al., 2009), and brainstem (Mena-Segovia et al., 2008) are locked to the phase of the slow cortical oscillation during slow wave sleep and anesthesia. Indeed, these results highlight the strong entrainment capacity of slow cortical oscillation ‘up’ states on downstream sub-cortical networks. However, local spindle-like activity in the SC was also locked to the phase of cortical slow waves, displaying a strikingly similar cross-frequency phase-amplitude relationship as thalamically generated spindles (Crunelli and Hughes, 2010; Steriade et al., 1993a, 1993b). Taken together, the entrainment of SC activity to thalamocortically coordinated sleep oscillations may suggest that the SC plays some role in regulation of neural activity during sleep (Miller et al., 1998), however further studies in chronically implanted animals would be needed to fully elucidate any sleep function of the SC.

### High frequency LFP components

Despite the transmembrane potential being one of the most informative neurophysiological signals in the brain, limitations of large-scale electrophysiological recording techniques have required that investigators infer intracellular processes from the less informative extracellular field, as measured by LFPs. The reason that there has been an expanding interest in higher frequency components of extracellular fields is that these signals have been shown to track the spiking activity neurons (Buzsáki et al., 2012; Manning et al., 2009; Ray and Maunsell, 2011), and the trial-to-trial behavioral performance of ECoG-implanted epilepsy patients (Szczepanski et al., 2014). Indeed, a recent study by Khodagholy et al (2014) showed that the spiking activity of individual superficial cortical neurons can be detected by recording cortical surface potentials with sufficiently small electrodes (10 x 10 μm^2^). In addition to reflecting Na^+^ action potentials, high frequency cortical surface potentials result from the superposition of several ionic processes that coincide with neuronal firing, such as synaptic currents, calcium spikes, spike afterhyperpolarizations, and intrinsic currents and resonances (Buzsáki et al., 2012; Trevelyan, 2009). Thus, although spiking activity correlates with high frequency liECoG activity, the relative contribution of the abovementioned ionic processes to the composition of high frequency LFP signals reported in this study remains unclear. Nevertheless, our results extend on previous work by illustrating that coincident fluctuations in high frequency signals between areas reflect the synchronized spiking activity of widely distributed neuron populations. Indeed, this finding has strong implications for human ECoG studies in particular, by enabling the inference of spiking processes without penetrating the cortical surface.

In conclusion, we present a novel approach for investigating large-scale cortico-subcortical functional interaction through simultaneous depth and μECoG recordings. This approach enabled us to identify multiple ongoing cortico-tectal coupling modes spanning different carrier frequency bands. High frequency power envelope correlations conveyed the most information about direct cortex-to-SC communication, and acted as a proxy for detecting coordinated spiking activity. Our results speak towards scaling down the dimensions of ECoG grids used in human studies in favor of high-density and small diameter electrode designs, which would enable the tracking of spiking activity with high spatial resolution over vast areas of cortex and improve the accuracy of detecting pathologically altered neural activity and abnormal modes of neural communication.

## Experimental procedures

Data presented in this study were collected from nine adult female ferrets *(Mustela putorius)*. All experiments were approved by the Hamburg state authority for animal welfare (BUG-Hamburg) and were performed in accordance with the guidelines of the German Animal Protection Law.

**Custom μECoG design:** For large-scale recordings of cortical LFPs, we used a polyimide based micro-electrocorticographic electrode array (μECoG) (Rubehn et al., 2009), with a thickness of 10 μm which was developed to optimally fit to the posterior cortex of the ferret. The custom μECoG array consisted of three ‘fingers’, each containing three rows of electrodes such that the polyimide foil can bend and conform to the curved surface of the ferret brain. The μECoG array had 64 platinum thin-film electrodes with a diameter of 250μm. Electrodes were arranged in a hexagonal formation with an interelectrode distance of 1.5mm. Figure 1A displays a schematic diagram of the μECoG layout. To allow for simultaneous recording of μECoG signals and intracortical activity, small holes of 500 μm diameter were cut into the polyimide foil in the space between electrodes to allow for the placement of multichannel linear probes.

**Surgery:** Animals were initially anesthetized with an injection of ketamine (15mg/kg). A glass tube was placed in the trachea to allow artificial ventilation of the animal and supply isoflurane anesthesia (0.5-1%, 1:1 NO - O_2_ mix). To prevent dehydration of the animal throughout experiments, a cannula was inserted into the femoral vein to deliver a continuous infusion of 0.9% NaCl, 0.5% NaHCO_3_ and pancuronium bromide (60μg/kg/h). Physiological parameters such as the ECG, rectal temperature and end tidal CO_2_ concentration were monitored throughout experiments to maintain the state o the animal. A large craniotomy was then performed over the entire left posterior cortex. After carefully removing the dura, the μECoG array was gently placed on the surface of the cortex. A small hole was drilled in the removed piece of scull over the area corresponding to the visual cortex, and then the piece of bone was fixed back in place with silicone elastomer (World Precision Instruments). All recordings were carried out in a dark sound attenuated chamber.

**Electrophysiology:** Neural activity in the SC was obtained with 2x16 channel dual-shank silicon probes (NeuroNexus Technologies, 100μm electrode spacing, 500μm inter-shank distance). Intracortical activity was obtained with linear 1x32 channel (100μm electrode spacing, NeuroNexus Technologies) probes that were inserted into the visual cortex through small holes in the μECoG (Figure 1). All silicon probe contacts had a surface area of 413μm^2^ . Broadband signals from silicon probes were digitized at 22321.4Hz (0.1Hz high pass and 6000Hz low pass filters), while μECoG signals were digitized at 1395.1Hz (0.1Hz high pass and 357Hz low pass filters). Broadband data from silicon probes and μECoG were sampled simultaneously with a 128 channel AlphaLab SnR^TM^ recording system (Alpha Omega Engineering). Unless otherwise stated, spontaneous recordings of neural activity of approximately 10 minutes in length were used for functional connectivity analyses presented in this paper.

**Visual stimulation:** Visual stimuli consisted of flashes and drifting gratings presented on an LCD monitor (Samsung SyncMaster 2233, 60Hz refresh rate) placed 28.5cm in front of the animal. Stimuli were generated using the Matlab Psychophysics Toolbox (The Mathworks Inc, MA). Very large (40°) flashes were used for probing visual responses, while smaller flashes (8°) were used for quantifying visual spatial receptive fields. To provide more sustained visual stimulation, drifting gratings of 1 second duration were presented in 12 different directions spanning 360°.

**Alignment of μECoG and SC electrode placement across animals:** For each animal, the position of the μECoG over the posterior cortex was recorded by taking photographs through a Zeiss OPMI pico microscope. The approximate position of all 64 μECoG electrodes was then projected onto a scaled illustration of a model ferret brain. The cortical topography of power correlation measures was then interpolated to fill the space between μECoG electrodes. Finally, cortical topographies for all SC-μECoG recording penetrations were averaged to obtain the mean power correlation topography. SC penetrations were aligned using the flash current source density inflection depth (Stitt et al., 2013). For more details on SC-depth realignment, see Supplementary Figure 2.

**Data Analysis:** All offline data analysis of neural signals was performed using custom software in Matlab (The Mathworks Inc, MA). To extract multiunit spiking activity (MUA) from broadband extracellular recordings, we high-pass filtered signals at 500Hz and detected spikes with a positive and negative threshold (Quiroga et al., 2004). LFPs were obtained by low-pass filtering broadband signals at 300 Hz. LFPs were filtered in both the forward and reverse direction to ensure zero phase shift. Finally, LFPs were downsampled by a factor of 16 to a sample rate of 1395.1 Hz. We computed spectral estimates of LFPs using a series of 80 Morlet wavelets that were logarithmically spaced from 0.1 – 300 Hz (Fieldtrip, width = 7 cycles), with increments of 5 ms.

*Power correlation*: To track the waxing and waning of neural signals over time, we computed time-resolved estimates of the LFP power. To avoid the detection of spurious power correlation due to the effects of shared noise or volume conduction, we orthogonalized pairs of signals prior to the computation of power envelope time series (Hipp et al., 2012). This crucial step removes zero-phase lagged components shared between simultaneously recorded signals, and ensures that only non-zero phase-lagged signal components are considered for power envelope correlation analysis. Following orthogonalization, power envelopes we computed by taking the absolute value of the square of complex Fourier spectra. The strength of power correlation was then defined as the linear correlation coefficient of orthogonalized power envelope time series. Significant SC-μECoG power correlation channel pairs were defined as those that exceeded the mean plus two times the standard deviation of the global SC-μECoG power correlation matrix (p < 0.05).

*Spike-phase locking:* To estimate the influence of the phase of cortical oscillations on spiking activity in the SC, we calculated spike-LFP phase-locking values (PLV) using spikes recorded from the SC and the phase of μECoG signals (Lachaux et al., 1999). For each SC-μECoG channel pair, spike PLVs were calculated from 1000 randomly drawn spikes. This process was repeated 100 times to get an estimate of the mean spike PLV. Channels where less than 1000 spikes were detected in the entire recording were eliminated from spike phase locking analysis. The level of significance for spike PLVs was determined by the Rayleigh statistic (Fisher, 1993).

*Cross-frequency coupling:* We quantified cross-frequency phase-amplitude coupling using methods previously described by Canolty et al (2006). Briefly, for each SC-μECoG channel pair, the phase of cortical signals, and the amplitude of SC signals were combined to construct composite Fourier spectra. The magnitude of the mean of composite signals provides an indication of the strength of phase-amplitude coupling. To estimate the standard deviation of cross-frequency coupling measures, this analysis was repeated 20 times with randomly shifted SC-amplitude time series. Finally, the modulation index was defined as the mean of composite spectra expressed in terms of the standard deviation of randomly shifted data.

## Acknowledgements

We would like to thank Dorrit Bystron for assistance throughout experiments. This research was supported by funding from the DFG (GRK 1246/1-2; SFB 936/A2; SPP 1665/EN/533/13-1; A.K.E.).

---

**Figure S1.**
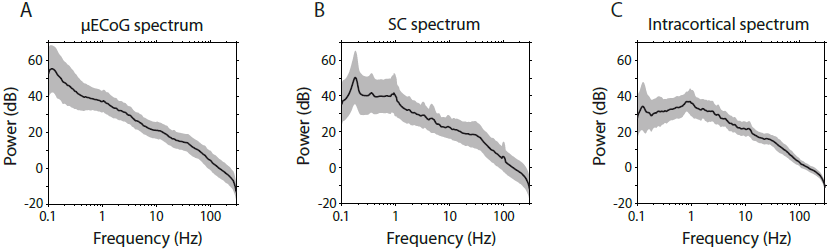
Panels **A-C** display the population average power spectra of μECoG, SC and intracortical recording sites respectively (± standard error mean).

**Figure S2.**
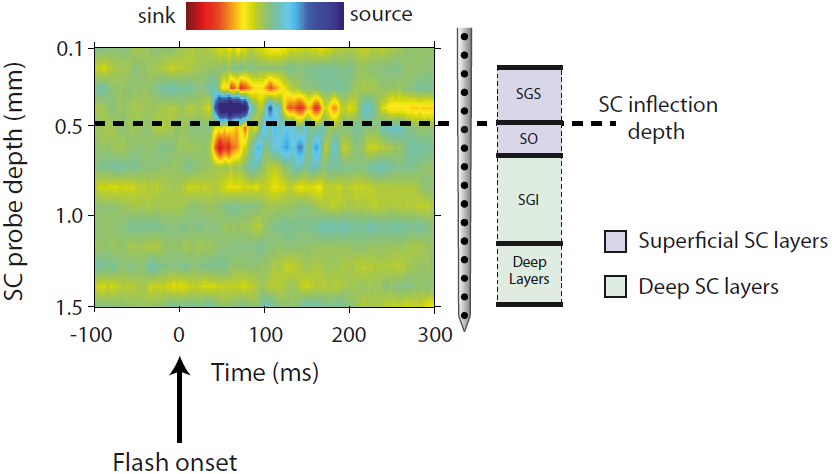
Recording depth in the SC was determined by current source density analysis. In a previous study (Stitt et al., 2013), our lab demonstrated that the border between flash-evoked current sources and sinks corresponds to the anatomical border between the two main superficial SC layers, the SGS and SO. Recording contacts were separated into two groups for depth-wise analysis, superficial SC – encompassing channels 300μm above and 200μm below the inflection depth, and deep SC – all channels situated deeper than 200μm below the inflection depth. Abbreviations: SGS – *stratum grieum superficiale;* SO – *stratum opticum*; SGI – *stratum griseum intermediale*.

**Figure S3.**
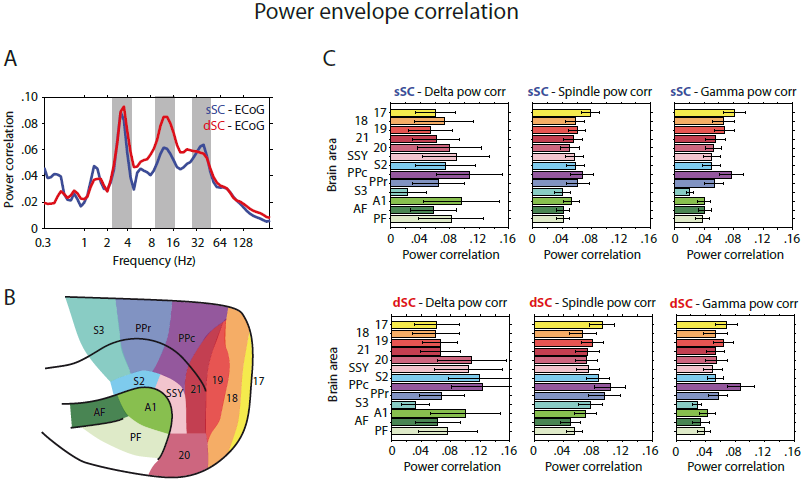
Large-scale structure of envelope coupling modes. **A:** Population averaged cortico-tectal power envelope correlation computed for SC recording contacts located in superficial (blue) and deep (red) SC layers. Note the peaks in cortico-tectal power correlation for delta, spindle and gamma frequencies. **B:** Displays the parcellation of the cortical surface into 13 different anatomically and functionally specialized regions. **C:** Displays the population averaged (± SEM), region-specific strength of cortico-tectal power correlation the delta, spindle, and gamma frequencies identified in **A.** Superficial SC-μECoG channel pairs are plotted in the top row, with deep SC-μECoG channel pairs plotted below. Abbreviations: SSY – suprasylvian area; PPc – posterior parietal caudal area; PPr – posterior parietal rostral area; S2 – second somatosensory area; S3 – third somatosensory area; A1 – primary auditory cortex; AF – anterior auditory areas; PF – posterior auditory areas.

**Figure S4.**
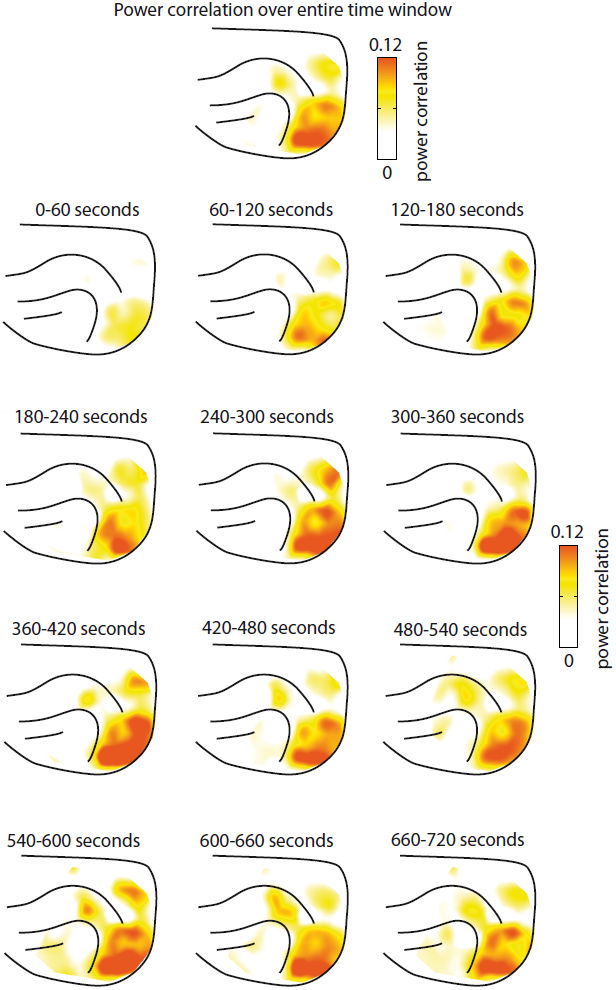
Temporal variability of cortico-tectal high frequency power envelope correlation. **A:** An example cortical topography of SC-μECoG power correlation computed over approximately 12 minutes of data. **B:** Displays the strength of cortico-tectal power correlation for the same example displayed in **A**, with correlation computed for 12 separate nonoverlapping one-minute blocks. Note that the strength of power correlation displays variation over time, however the topography remains constant.

**Figure S5.**
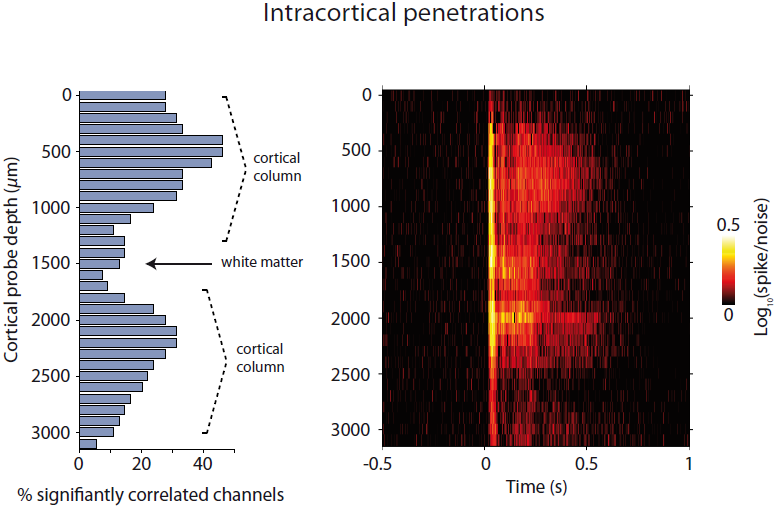
Cortical depth profile. **A:** Displays the incidence of channels that are significantly correlated with the SC and a function of cortical depth. Note that power correlation effects display two peaks at 500μm and 2200μm below the cortical surface respectively, reflecting that we usually recorded form two vertically adjacent cortical penetrations separated by white matter. **B:** Displays the population averaged spike/noise ratio for cortical spiking activity in response to visual flash stimulation.

**Figure S6.**
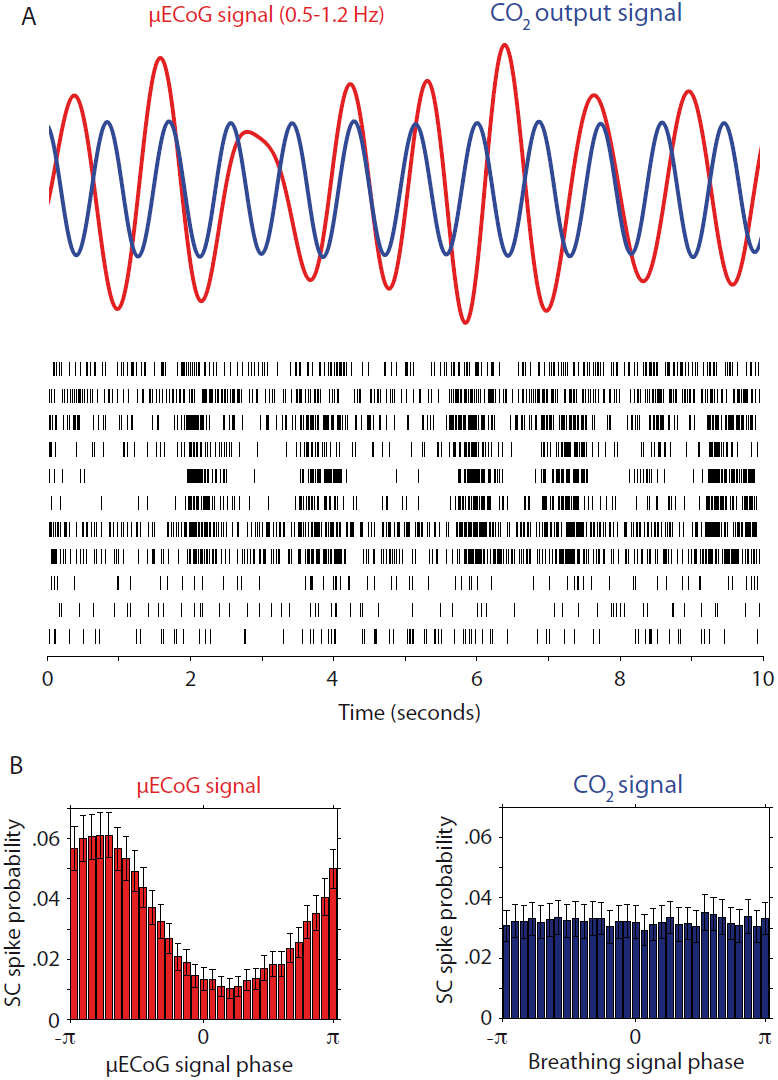
Effects of breathing on coordinated neural activity. **A:** Displays a filtered μECoG signal time-series plotted along with the simultaneously recorded output of the carbon dioxide breathing monitor. The breathing monitor indirectly tracks any mechanical related brain pulsations caused by ventilation. Note that the endogenous slow brain oscillation and breathing signal appear entirely unrelated. **B:** Displays spike-phase histograms computed using SC spiking activity and the phase of μECoG (left) and breathing (right) signals. Note that SC spiking activity is strongly locked to the phase of the endogenous physiological oscillation, and displays no temporal dependency on breathing.

**Figure S7.**
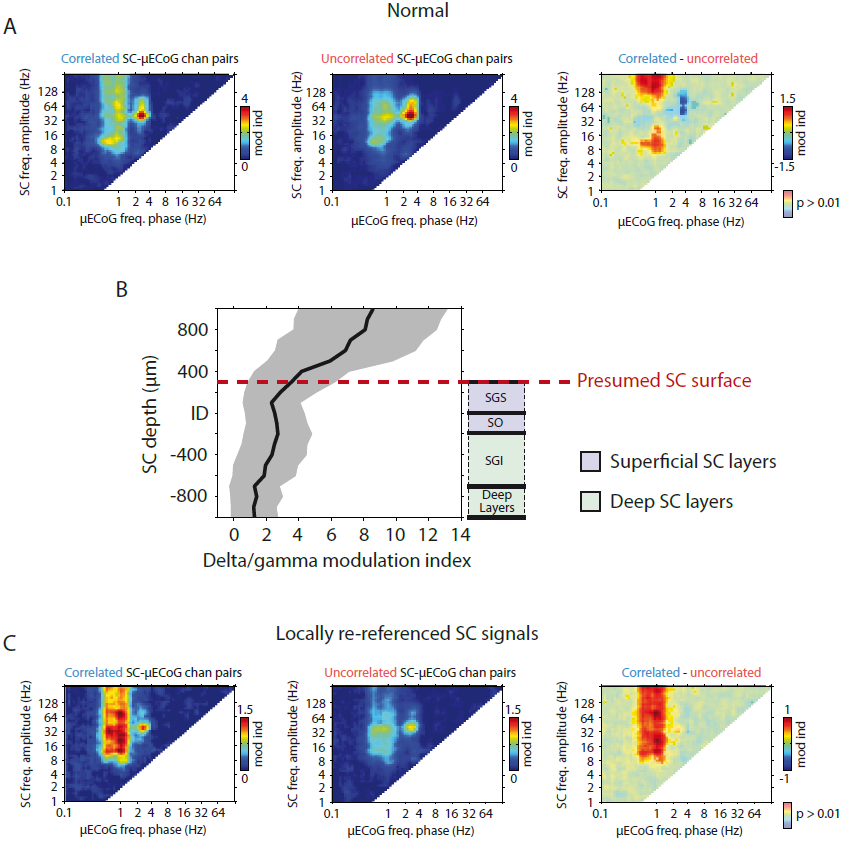
Cross frequency coupling spectra and locally re-referenced signals. **A:** Displays the population averaged cross-frequency phase amplitude coupling spectra for power correlated (left) and uncorrelated (middle) SC-μECoG channel pairs. The difference between correlated and uncorrelated channel pairs is plotted on the right (p < 0.01, Bonferroni corrected). Note the significant locking of spindle and high frequency activity to the phase of slow cortical oscillations. In addition, both correlated and uncorrelated channel pairs display very strong cross-frequency coupling between cortical delta phase and SC gamma activity. **B:** Displays the population average strength of cortico-tectal delta/gamma phase amplitude coupling as a function of SC depth. Note that the strength of delta/gamma coupling drastically increases for recording contacts that are presumed to be located superficial to the SC. This result suggests that observed delta/gamma coupling may in fact be due to volume conduction from the brain structure directly overlying the SC. **C:** Displays the same analysis as performed in A, however following the local re-referencing of SC signals. Note that the strong delta/gamma cross-frequency coupling present in **A** is now weakened or absent.

